# Resilience and Charge-Dependent Fibrillation of Functional Amyloids: Interactions of *Pseudomonas* Biofilm-Associated FapB and FapC

**DOI:** 10.1101/2024.02.14.580233

**Authors:** Nimrod Golan, Amit Parizat, Orly Tabachnikov, Eilon Barnea, William P. Olsen, Daniel E. Otzen, Meytal Landau

## Abstract

FapC and FapB are biofilm-associated amyloids involved in the virulence of *Pseudomonas* and other bacteria. We herein demonstrate their exceptional thermal and chemical resilience, suggesting that biofilm structures might withstand standard sterilization, thereby contributing to the persistence of *P. aeruginosa* infections. Our findings also underscore the impact of environmental factors on Fap proteins, suggesting that orthologs in different *Pseudomonas* strains adapt to specific environments and roles. Challenging previous assumptions about a simple nucleation role for FapB in promoting FapC aggregation, the study shows a significant influence of FapC on FapB aggregation. The interaction between FapB and FapC is intricate: FapB stabilizes FapC fibrils, while FapC slows down FapB fibrillation but can still serve as a cross-seeding template. This complex interplay is key to understanding their roles in bacterial biofilms. Furthermore, the study highlights distinct differences between Fap and *E. coli*’s curli CsgA amyloid, where CsgB assumes a simple unidirectional role in nucleating CsgA fibrillation, emphasizing the importance of a comprehensive understanding of various amyloid systems. This knowledge is vital for developing effective intervention strategies against bacterial infections and leveraging the unique properties of these amyloids in technological applications such as novel bio-nanomaterials or protective coatings.

## Introduction

The opportunistic pathogen *Pseudomonas aeruginosa* is a significant cause of mortality in patients with co-morbidities, particularly those with cystic fibrosis (CF) and ventilator-associated pneumonia^1–5^ . The resistance of this bacterium to various antibiotics underscores the urgent need for novel treatment approaches^1,4–7^. In CF patients, chronic respiratory infections caused by *P. aeruginosa* typically begin in adolescence and can persist for decades^2,3^. The bacterium adapts to the pulmonary environment by forming biofilms that adhere to the respiratory epithelium, thereby evading the immune response and shedding immunogenic structures like pili and flagella^1–3^.

Biofilms, which enhance surface adherence and create a protective environment for bacterial growth, are observed in *Pseudomonas* strains across diverse environments^8–10^. In 2010, Dueholm *et al.* provided evidence of amyloid fibril involvement in *Pseudomonas* biofilm formation^11^. Amyloids are known for their remarkable stability, resistance to protease digestion and denaturation, and self-polymerizing capabilities, serving as robust structural elements^4,12–14^. In microbes, amyloids can serve as scaffolds in biofilm matrices, especially under harsh and energy-depleted conditions^4,8–10,15^. Amyloid fibrils may also act as surfactants and adhesins and are thought to play a role in bacterial quorum sensing systems ^16,17^.

Bacterial amyloids are regulated by specific pathways, beneficial for the physiology of the organism, frequently aided by an assortment of accessory proteins^18^. In *Pseudomonas*, amyloid production is orchestrated by the six-gene operon named Functional amyloid in *Pseudomonas* (Fap), with FapC and FapB serving as the amyloid-forming subunits^4,8,11^. In enterobacteria, the curli system is tightly controlled by two operons, CsgBAC and CsgDEFG, with CsgA as the primary amyloid subunit and CsgB as a nucleator of CsgA fibrillation^19^. Drawing parallels with the curli system, it was hypothesized that Faps serve similar roles as CsgA/B^8–11,20–22^. Accordingly, FapB could function as a nucleator, accelerating the rapid elongation of FapC fibrils on the cell surface, or it might be integral to the structure of the mature fibril, thereby altering its physicochemical characteristics^4,11,19^. FapA influences the distribution of FapC and FapB in mature fibrils, with its deletion leading to FapB-dominated biofilm fibrils^8,9,11,20,23^. FapD, a putative cysteine protease, may initiate secretion before degradation^24^, while FapE’s role as an extracellular chaperone is still being explored. The membrane protein FapF forms trimeric β-barrel channels, as observed by a crystal structure^24^, crucial for extracellular Fap component secretion^4,8,23,24^. Fap proteins display various virulence traits. Removing FapC in *P. aeruginosa* reduces virulence, as shown in *Caenorhabditis elegans* models and polymorphonuclear neutrophil leukocytes phagocytosis assays^25,26^. The Fap system is not exclusive to *P. aeruginosa*; other pathogenic bacteria also possess Fap-encoding genes, like *Aeromonas caviae*, and *Laribacter hongkongensis*, known to cause gastroenteritis and diarrhea, as well as *Burkholderia gladioli*, *B. pseudomallei*, *Ralstonia pikettii*, and *Stenotrophomonas maltophilia*, which are associated with airway and lung infections^25,27–30^.

Advances in structural biology have shed light on the stability and complex nature of eukaryotic amyloids. The prominent structural feature of amyloids is the formation of cross-β fibrils comprising paired β-sheets made of β-strands aligned perpendicularly to the fibril growth axis^4,13,31,32^. In contrast to human amyloids, understanding bacterial amyloids has been hampered by a lack of high-resolution structural data. A breakthrough came with the first crystal structure of bacterial amyloid, the cytotoxic PSMα3 from *Staphylococcus aureus*, revealing a polymorphic amyloid fold where α-helices can replace β-strands to form cross-α fibrils, contributing to its pathogenicity^33^. Recent advancements in cryogenic electron microscopy (cryo-EM) have begun to unveil the structures of biofilm-associated amyloids, highlighting both similarities and distinctions when compared to other amyloids found in humans and microbes. The cryo-EM structure of TasA from *Bacillus subtilis* consists of a fiber formed from folded monomers. These monomers are assembled through a donor–strand exchange mechanism, where each unit contributes a β-strand to complete the fold of the subsequent subunit in the fiber^34^. In contrast, the structure of *E. coli* curli CsgA, elucidated using a synergistic approach of CryoEM and computational modeling, exhibited a unique globular β-solenoid configuration, composed of monomeric units that enable a quasi-homotypic stacking of β-strands^35^. The predicted structures of FapC, FapB and FapE modeled using AlphaFold^36,37^ suggest a β-solenoid which resembles the structure of CsgA^38^. The structural insights into Faps, CsgA and TasA not only challenge traditional views of protein polymers and amyloid fibrils but also suggest that our understanding of functional amyloids may need reevaluation and expansion to encompass novel structural paradigms.

Here we delve into the molecular interactions of *P. aeruginosa* FapB and FapC, focusing on their fibrillation behavior and interaction dynamics. We explore their fibrillation rates, pH sensitivity, charge distribution, and response to various conditions, including their thermostability and chemical resistance. Our findings argue against the role of FapB as a nucleator, at least in the PAO1 strain, and rather suggest a greater effect of FapC on FapB in manipulating fibrillation rate and forming co-localized aggregates. Moreover, our findings on their resilience to extreme conditions, such as high temperatures and formic acid, highlight the challenges they pose in medical settings due to their resistance to standard sterilization methods. Overall, our study not only advances the understanding of bacterial amyloids dynamic nature but also emphasizes the necessity for innovative strategies to tackle biofilm-associated infections in *P. aeruginosa*.

## Results

### Differential sensitivity of FapB and FapC to pH and ionic strength conditions

In our study, we explored the differential sensitivity of FapB and FapC from the *Pseudomonas aeruginosa* PAO1 strain (UniProt IDs Q9I2E9 and Q9I2F0, respectively) to variations in pH and ionic strength. We expressed and purified these proteins and assessed their fibrillation rates using a Thioflavin T (ThT) fluorescence assay across a range of pH levels (Figure 1), different salt concentrations (Figure S1), tag location (Figure S2), and different protein concentrations (Figure S3).

**Figure 1:**
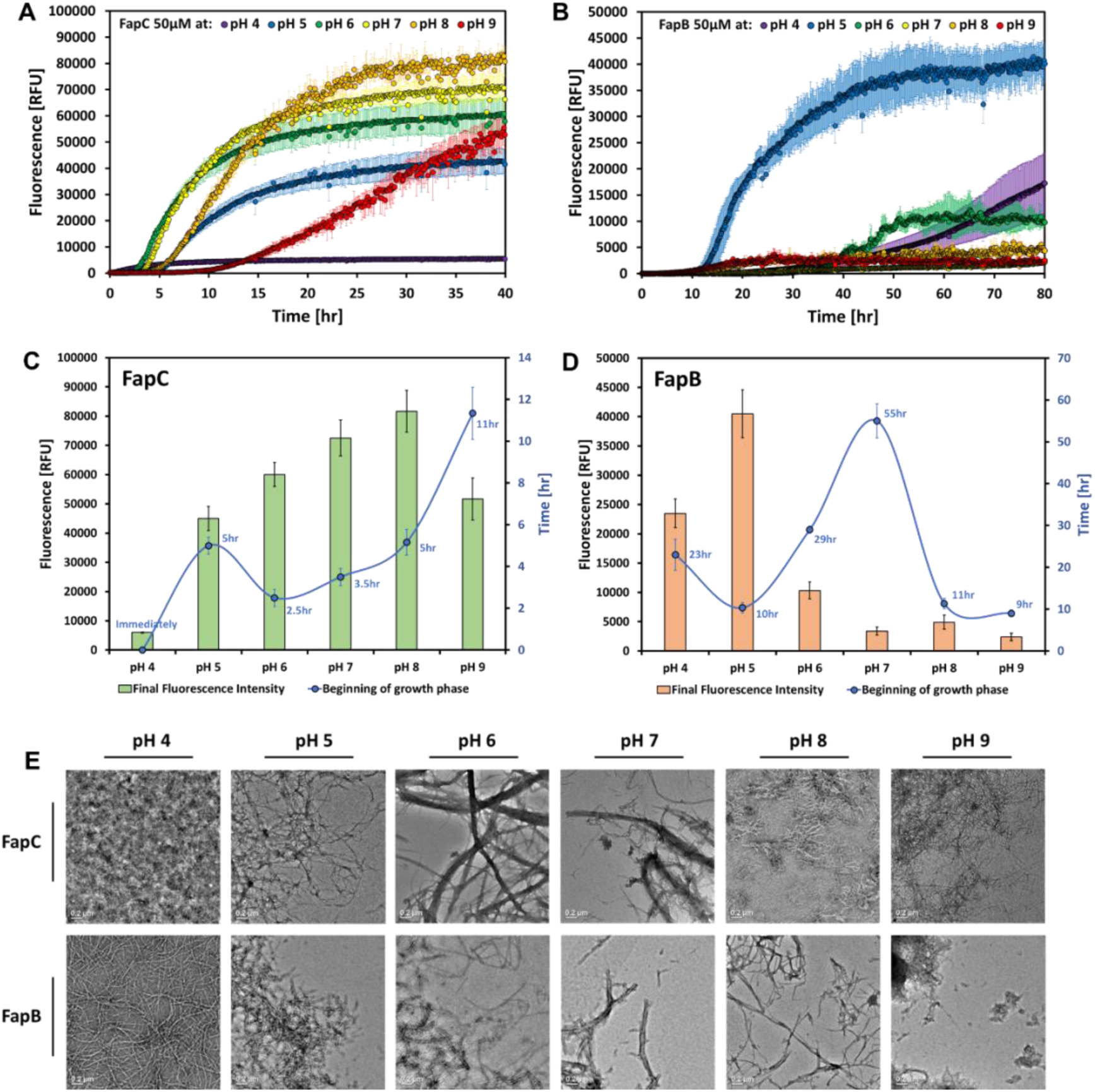
Influence of pH on the fibrillation rates of FapB and FapC. ThT was used to monitor fibrillation of 50 μM FapC (A) and FapB (B) between pH 4 and 9. The error bars in the graphs represent the standard deviation from the average of triplicates. The experiments were carried out on three separate days with consistent results. For enhanced clarity, the graphs detail the quantification of the onset of the growth phase (lag time) and the final fluorescence intensity for both FapC (C) and FapB (D) under various pH conditions. (E) Transmission electron micrographs showcase the morphology of 50 μM FapB and FapC fibrils after 80 hours of incubation under six pH conditions, highlighting the influence of pH on fibril morphology. All micrographs include a 200 nm scale bar.

FapC, at a concentration of 50 μM, displayed a characteristic amyloid fibrillation profile, characterized by a lag phase, a growth phase, and typical sigmoidal fibrillation kinetics (Figure 1A). Optimal fibrillation for FapC, with a lag time of approximately three hours, was observed at neutral pH values (6-7). The rate decreased slightly at pH 5 and pH 8, with a more pronounced reduction at pH 9 (Figure 1C). Notably, at pH 4, the fibrillation curve remained almost flat over a 40-hour incubation period (Figure 1A). Altering the ionic strength by varying NaCl concentrations (0 M, 0.15 M, and 0.3 M) at pH 7 did not significantly impact the fibrillation curves of FapC (Figure S1).

The fibrillation rate of 50 μM FapB was significantly influenced by pH, exhibiting the fastest rate (lag time of about 10 hours) at pH 5 (Figure 1D). Weakly acidic conditions (pH 4-6) were generally most conducive for FapB fibrillation (Figure 1B). Higher NaCl concentrations marginally increased the slope in ThT time curves but did not alter the lag time (Figure S1). Notably, the position of the Histidine tag on FapB affected its fibrillation rate. The C-terminal Histidine tag almost completely blocked fibrillation of FapB at pH values between 5 and 9 (Figure S2A). Consequently, all subsequent experiments were conducted using N-terminus Histidine tagged FapB and C-terminus Histidine tagged FapC.

Transmission electron microscopy (TEM) analyses after 80 hours of incubation showed the formation of fibrils for both FapB and FapC under almost all pH conditions, though with differences in abundance and morphology (Figure 1E). A notable distinction was observed at pH 4, where FapB formed thin, long, and well-separated fibrils, while FapC primarily produced dense bundles of aggregates where the fibrillar structure could barely be seen. In contrast, at the other extreme case of pH 9, FapC fibrils were more prevalent, indicating a preference for fibrillation in neutral to basic conditions for FapC and in acidic conditions for FapB.

Building on the established optimal pH conditions for FapB (pH 5) and FapC (pH 7), we further investigated their fibrillation rates at additional concentrations (Figure S3). FapC showed a dose-dependent increase in both fibrillation rate and ThT fluorescence intensity at both pH levels, with a lag time at 50 μM shorter than 10 hours. In distinction to FapC, FapB demonstrated a rapid fibrillation rate of 5-12 hours only at pH 5, with a dose-dependent reduction in lag time and an enhanced ThT fluorescence signal. However, at pH 7, FapB’s fibrillation rate was considerably slower, exceeding 25 hours, and the kinetic curve deviated from the typical sigmoidal shape, not reaching a plateau even after 70 hours.

These observations underscore the distinct pH-dependent fibrillation behaviors of FapB and FapC, highlighting their unique responses to environmental conditions and suggesting potential functional differences in biofilm formation and stability.

### Net charge and pH sensitivity in Fap orthologs

FapB and FapC from various *Pseudomonas* strains, including *P. aeruginosa* PAO1, *P. fluorescens* Pf-5, *P. putida* F1, and *Pseudomonas sp*. UK4, exhibit significant variations in length, charge, and isoelectric point, as detailed in Table S1 along with their Uniprot IDs. FapC and FapB are structured with three imperfect sequence repeats, linked together by two regions.

Additionally, structural modeling using AlphaFold indicates that these proteins adopt a Greek Key β-solenoid configuration^38^. The shorter linkers in FapB result in comparatively shorter sequences, likely contributing to smaller length variations among its orthologs. The sequence lengths of FapC homologs show considerable diversity, ranging from 226 amino acids in *Pseudomonas sp*. UK4 to 459 in *P. putida* F1. In contrast, the lengths of FapB homologs across these strains are more uniform, with a range from 164 amino acids in *Pseudomonas sp.* UK4 to 174 in *P. putida* F1. Despite these variations in length, FapB and FapC in the PAO1 *P. aeruginosa* strain we studied here, which are 170 and 316 residues long respectively, share a 38% sequence identity^4^. This contrasts with *E. coli* CsgA and CsgB, which, despite having similar lengths (131 and 130 residues respectively), share only 23% sequence identity.

FapB and FapC and their orthologs exhibit distinct variations in their charge distribution and sensitivity to pH. FapC orthologs, despite their different lengths, showed a narrow spectrum from slightly acidic to neutral isoelectric points, yet exhibited a broad range of total charge at neutral pH, from -14.02 to -0.13 (Table S1). The total charges of FapC orthologs were also significantly affected by pH changes (Figure 2A). For example, *P. putida* F1’s FapC net charge fluctuates from +20 at pH 4 to nearly -20 at pH 9, with an isoelectric point at pH 5.1 (Figure 2). FapC from *P. aeruginosa* PAO1 transitioned from a positive to a negative total charge above pH 4.9. With the addition of a Histidine tag at the C-terminus, as used in the study, this transition shifts to a higher pH of 5.7, yet the positive charge is mostly localized at the tag. Considering its rapid fibrillation over a wide pH range (5-8), as evidenced by a lag time of under 5 hours at 50 µM (Figure 1), this suggests that a negative charge may promote FapC fibrillation. However, at pH 9, the net charge could become overly negative, leading to repulsion that impedes orderly self-assembly. At pH 4, with a positive net charge, amorphous aggregates are formed instead of fibrils (Figure 1C).

**Figure 2:**
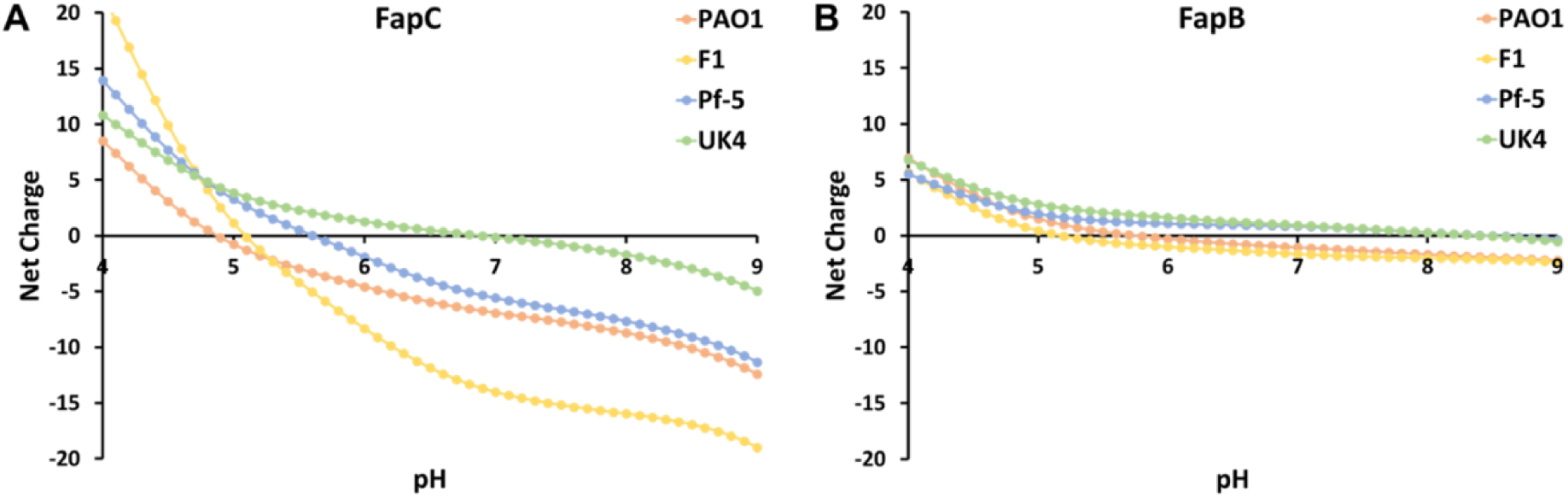
Comparative Analysis of Net Calculated Charge for FapB and FapC Orthologs. This figure illustrates the net calculated charge of four FapC (A) and FapB (B) orthologs across a pH spectrum from 4 to 9, calculated using the Prot-pi: Protein Tool for Isoelectric Point and Net Charge Calculation (https://www.protpi.ch/).

Unlike FapC orthologs, the isoelectric points of FapB orthologs exhibit considerable variation, spanning from acidic to basic. Despite this range, the overall charge of FapB orthologs at neutral pH remains relatively stable, oscillating between -1.64 and 0.94, as detailed in Table S1. Similarly, their net charge maintains a relatively neutral profile within a pH spectrum of 5-9, as shown in Figure 2B. Specifically, for *P. aeruginosa* PAO1 FapB, the isoelectric point is established at pH 5.8. The addition of an N-terminal Histidine tag shifts this point to 7.4, marking a more pronounced shift than that observed with FapC. To elucidate the impact of the tag on charge distribution, we employed an AlphaFold-generated model ^36,37^. of FapB at varying pH levels, both with and without the tag at either end (Figure S2B). This model revealed a folded monomer structure characterized by a β-solenoid with β-sheets that form interfaces akin to cross-β fibril structures. Notably, the surface charge distribution of this monomer remained consistent across the modeled pH levels, aligning with the predicted net charge variations (Figure 2B). The experimental examination of a C-terminal Histidine tag, featuring a short linker, demonstrated its interaction with FapB’s surface as per the model. This interaction alters the charge distribution across different pH levels, potentially explaining the linker’s role in inhibiting fibrillation (Figure S2A). Conversely, the model suggests that the N-terminal tag, possessing a longer linker, does not directly interact with FapB’s surface, correlating with the observed fibrillation capability of FapB tagged at the N-terminus. The pH-induced charge alterations of the tag predominantly occur at the periphery of FapB due to the extended linker. FapB tagged at the N-terminus significantly promotes fibrillation at pH 5, exhibiting a lag phase of approximately 10 hours at 50 µM concentration, in stark contrast to the over 30 hours observed at other pH levels (Figure 1). At this acidic pH, FapB’s surface assumes a relatively neutral charge, with the linked tag carrying a positive charge.

When considering these observations in the context of pH-dependent fibrillation rates, it becomes clear that charge distribution plays a significant role in regulating the driving forces and rate of fibril formation and morphology for both FapB and FapC. The behavior of FapB orthologs suggests a different mechanism of aggregation compared to FapC, potentially driven by the balance of charges rather than the accumulation of a specific charge polarity.

These findings propose a sophisticated regulatory mechanism where the protein’s charge significantly influences fibril formation rate and structure. Fibrillation of FapB, sensitive to pH and prone to aggregation near its isoelectric point, akin to the fibrillation observed in the peptide hormone glucagon^39^, might serve as an indicator of environmental changes. In contrast, FapC, fibrillating over a broader pH range, could act as a resilient amyloid, guided by specific electrostatic interactions due to its predominantly negative charge. Such charge and pH sensitivities might represent adaptive strategies, enabling these proteins to function effectively in varied conditions within bacterial biofilms.

### Delayed fibrillation through co-incubation of FapB and FapC monomers

FapB was hypothesized to act as a nucleator for the primary amyloid subunit FapC, akin to the role of CsgB with CsgA^8,11,19,20,40^. To investigate their potential reciprocal effects on fibrillation, we introduced various concentrations of fresh Fap monomers (from 2 µM to 100 µM) into a solution containing 50 µM of their counterparts, maintaining the optimal pH for each (Figure S4A&B). Figure S4C&D compare the fibrillation kinetics of Fap monomers incubated alone and as mixtures in both pHs and highlights three specific molar ratios (1:1.5, 1:1, and 2:1), while Figure 3A&B showcases the results for a representative 1:1 molar ratio. The co-incubation of Fap monomers resulted in a dose-dependent inhibitory effect, notably prolonging the lag phase of their fibrillation.

**Figure 3:**
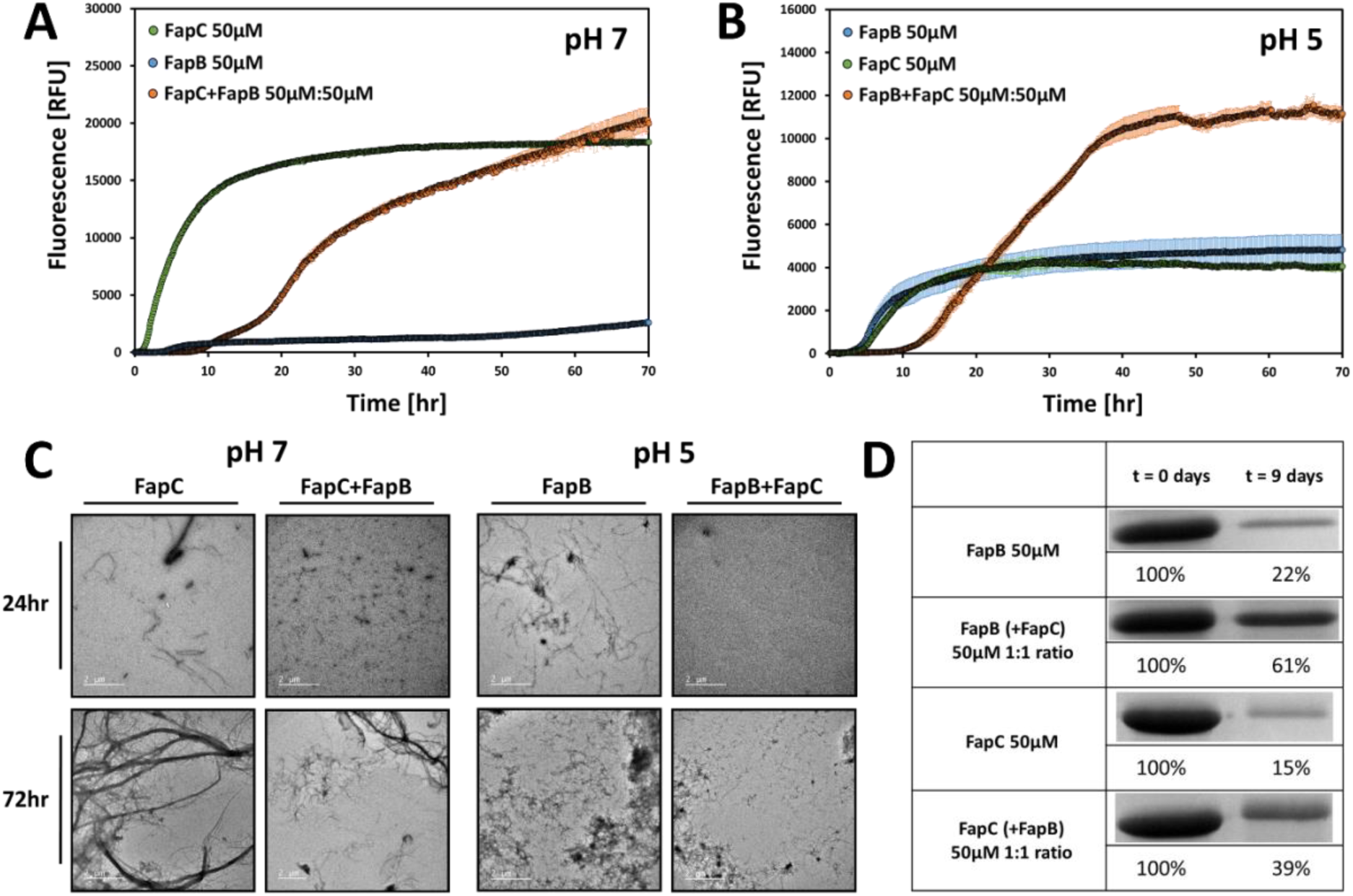
Impact of co-incubation on FapB and FapC fibrillation. ThT fluorescence kinetics showing the fibrillation rates of FapC at pH 7 (A) and FapB at pH 5 (B) both individually and when co-incubated with their respective counterparts at a 1:1 molar ratio (50:50μM). Error bars indicate standard deviation from three triplicate measurements. Repetitions of the experiment on different days yielded consistent results. The impact of varying molar ratios on fibrillation is detailed in Figure S4. (C) TEM images presenting 50 μM FapB (at pH 5) and FapC (at pH 7), incubated either separately or together in a 1:1 molar ratio for 24 and 72 hours. All images include a 2 μm scale bar for reference. (D) SDS-PAGE analysis illustrating the remaining amounts of FapB (at pH 5) and FapC (at pH 7) monomers after 9 days of incubation, both individually and in a mixture of 1:1 molar ratio. The bands obtained from the analysis were quantified, with each being represented as a percentage of the amount of monomers present in the pre-incubated sample.

The addition of FapB to FapC at pH 7 resulted in a dose-dependent extension of the lag phase compared to FapC fibrillation alone (Figures 3 & S4). This delay increased from a lag time of approximately 2 hours to as much as 15 hours with increasing concentrations of FapB. At 75 µM, FapB shows rapid fibrillation even at pH 7, with a lag time of approximately 15 hours, which seems to nevertheless halt the fibrillation rate of the 50 µM FapC (Figure S4C). This suggests that FapB, which fibrillates more slowly, reduces the fibrillation rate of FapC at a range of concentrations. At pH 5, a comparable dose-dependent inhibitory effect was observed, with the lag time of FapB extending from about 5 hours to as long as 25 hours following the addition of FapC (Figures S4B&D). Despite the significant fibrillation of FapC alone also at pH 5 (Figure S4D), its addition reduced the fibrillation rate of FapB (Figure 3B).

The bidirectional delay in fibrillation was visually confirmed by TEM images, which showed a reduced number of fibrils in the mixed samples after 24 hours of incubation at 50 µM 1:1 molar ratio at both pH levels. However, after 72 hours, this inhibition appeared to diminish, and fibrils formed in both mixtures, consistent with the kinetic assay results (Figure 3C). To quantitatively assess the extent of fibrillation, we exploited the resistance of Fap fibrils to sodium dodecyl sulfate (SDS) and boiling^8,11,41,42^ and measured the levels of residual monomers using SDS-PAGE before and after 9 days of incubation, both individually and in 1:1 mixtures (Figure 3D). The notable difference in molecular weight between the Fap proteins (Table S1) enabled precise detection of each individual Fap level. Analysis showed a higher monomer concentration in the mixed amyloid samples than in those where Faps were incubated separately, at both pH levels. Specifically, for FapC, after nine days, the monomer concentration was 15% of the pre-incubated sample when incubated alone, and 39% when mixed in a 1:1 ratio with FapB. In the case of FapB, the monomer concentration after nine days was 22% of the pre-incubated sample when incubated alone, and 61% when mixed in a 1:1 ratio with FapC. Overall, the data indicate that mixing different Fap proteins tends to decelerate their fibrillation process, with lasting effects even after nine days. The extent of this effect varies based on the specific Fap combination used, with FapC demonstrating a more pronounced influence on FapB.

### Self- and cross-seeding of FapB and FapC show minor but asymmetrical effects

To deepen our understanding of the interactions between FapB and FapC, we conducted self- and cross-seeding experiments using pre-formed, sonicated fibrils of these proteins. We assessed how these seeds influenced the fibrillation rates of 50 μM FapB or FapC monomers, by adding 2-10% (by total monomer sample volume) of FapB and FapC seeds (Figure 4). In both cases, self-seeding accelerated the fibrillation rate. This effect was especially pronounced for FapB, where it nearly eliminated the lag phase of fibrillation (Figure 4B). Cross-seeding experiments revealed that adding FapC seeds to FapB at pH 5 reduced the lag time in a dose-dependent manner (Figure 4A). However, when FapC was cross-seeded with FapB seeds, there was little to no effect on its fibrillation rate at any of the tested concentrations (Figure 4C).

**Figure 4:**
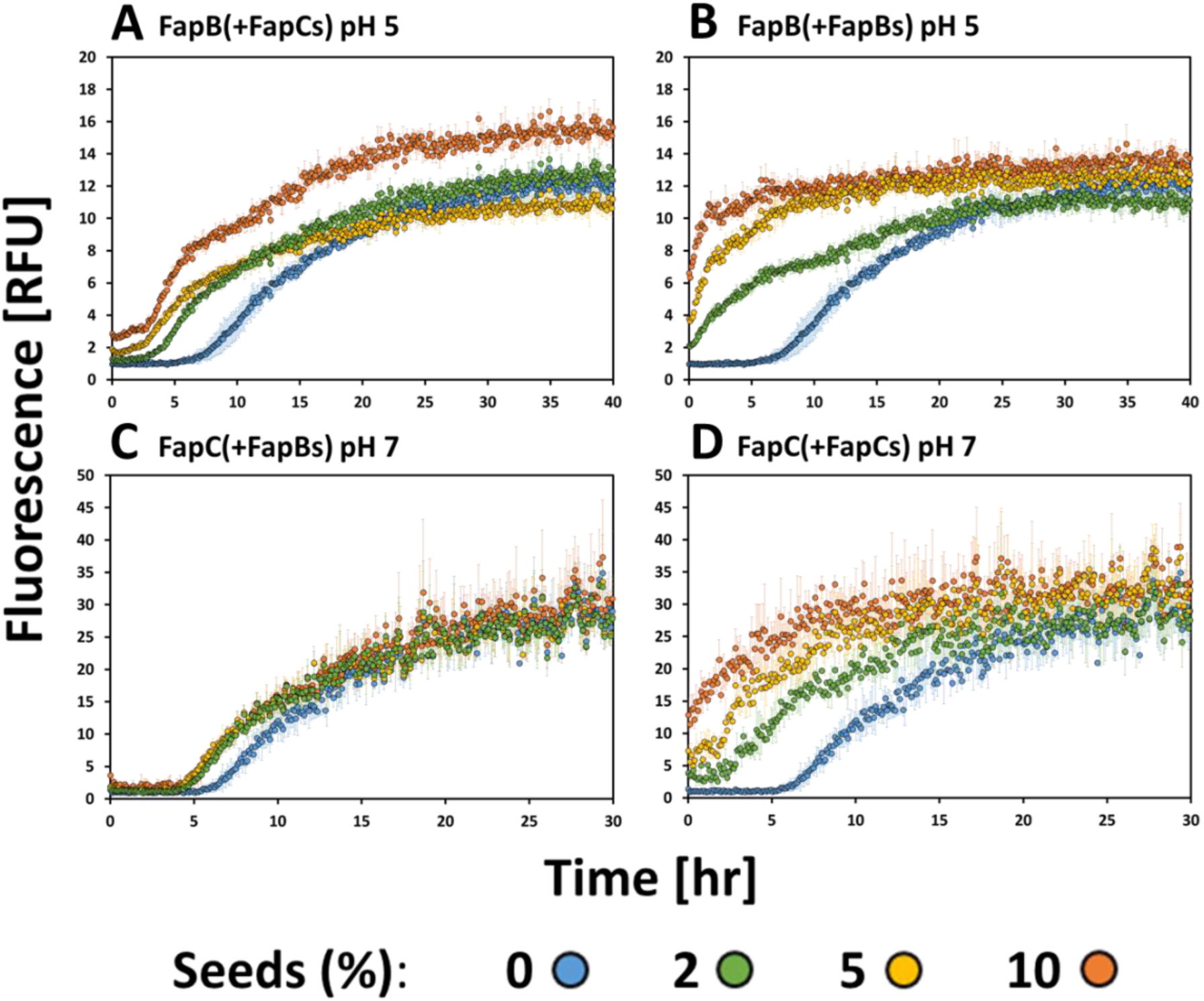
Effect of self- and cross-seeding on fibrillation rates of FapB and FapC. The figure demonstrates the impact of adding fibril seeds (indicated as “s” in the figure) of either FapB or FapC on the fibrillation rate of 50 μM FapB at pH 5 (panels A and B) and 50 μM FapC at pH 7 (panels C and D). Seeds were introduced at three different volume percentages of the total reaction mixture (2%, 5%, and 10%), and the fibrillation rate was monitored using ThT fluorescence. Error bars indicate the standard deviation from the mean of triplicate measurements. These experiments were replicated on separate days, consistently producing similar trends.

These results highlight an asymmetrical effect between the two Fap variants. FapC seeds had a more substantial influence on the fibrillation of FapB than the reverse, aligning with the patterns observed in monomer mixtures (Figure 3). This asymmetry suggests that FapB and FapC may follow different nucleation and elongation mechanisms in their amyloid formation and interactions. In addition, the observation that self-seeding had a more significant impact on accelerating fibrillation rates than cross-seeding, indicates distinct intra-molecular interactions for each Fap type. Although further research is needed to fully understand these mechanisms, the current findings suggest that FapB may not act as a nucleator for FapC, particularly in the tested orthologs from *P. aeruginosa* PAO1. Instead, FapB and FapC appear to have distinct roles in biofilm formation, adapting to various environmental conditions, and their interactions are more intricate than initially thought.

### Fluorescence microscopy reveals co-localization of FapB and FapC

Alterations in fibrillation rates observed during the co-incubation of Fap monomers point to a direct interaction between them. To further explore this, we employed confocal light microscopy for in-depth visualization. FapB and FapC were labeled with Cyanine3 (Cy3) and Cyanine5 (Cy5) fluorophores, respectively, using NHS ester crosslinking to attach these dyes to lysine residues within the proteins. The fibrillation of these labeled proteins was verified through TEM (Figure S5). Our analysis utilized two distinct approaches: first, we observed the aggregation and localization of fresh, monomeric Faps (Figure 5 and [Video 1](https://youtu.be/U-8s0u023T4)); second, we examined the interactions between preformed fibrils (Figure 6 and [Video 2](https://youtu.be/PPE1Kngsw3U)). Due to resolution constraints, the observed foci were referred to as aggregates, without distinguishing between fibrils and less ordered species.

**Figure 5.**
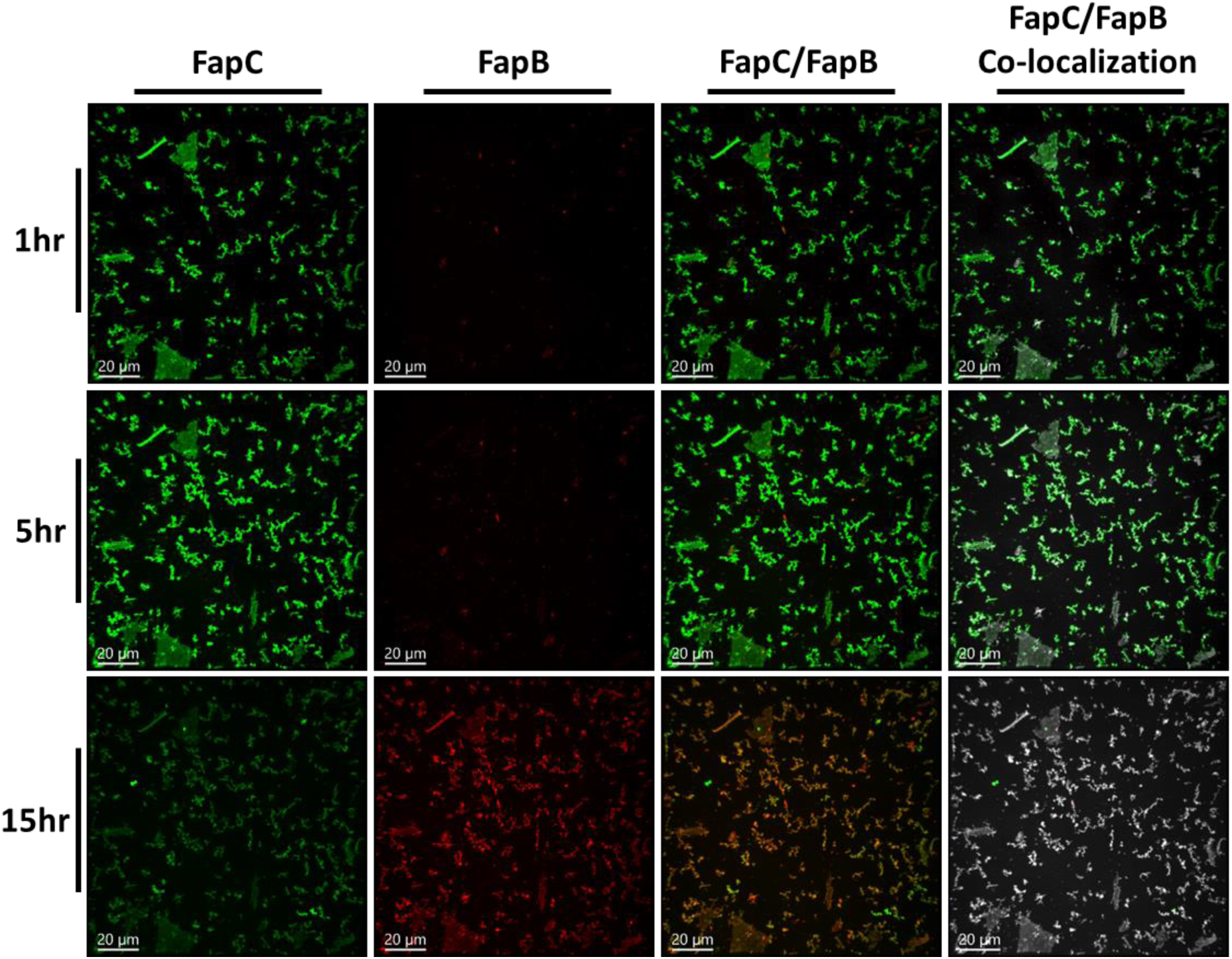
A dynamic visualization of FapB and FapC monomers co-localizing over time. Fluorescence microscopy images, captured from [Video 1](https://youtu.be/U-8s0u023T4), where FapB and FapC are distinctly labeled with Cy3 (red) and Cy5 (green) fluorophores, respectively, and co-incubated as fresh samples. The images capture the progression of aggregation and co-localization at 1, 5, and 15 hours, arranged in a top-to-bottom sequence. The merging of Cy3 and Cy5 channels results in a yellow/orange hue, pinpointing the areas where FapB and FapC aggregates intersect. For enhanced visual clarity, the right-hand side of each panel row illuminates these co-localization zones in white, as determined by computational analysis. Incubation during the time-lapse imaging was conducted at pH 7. Consistently included in all images are scale bars, each denoting a length of 20 μm.

**Figure 6.**
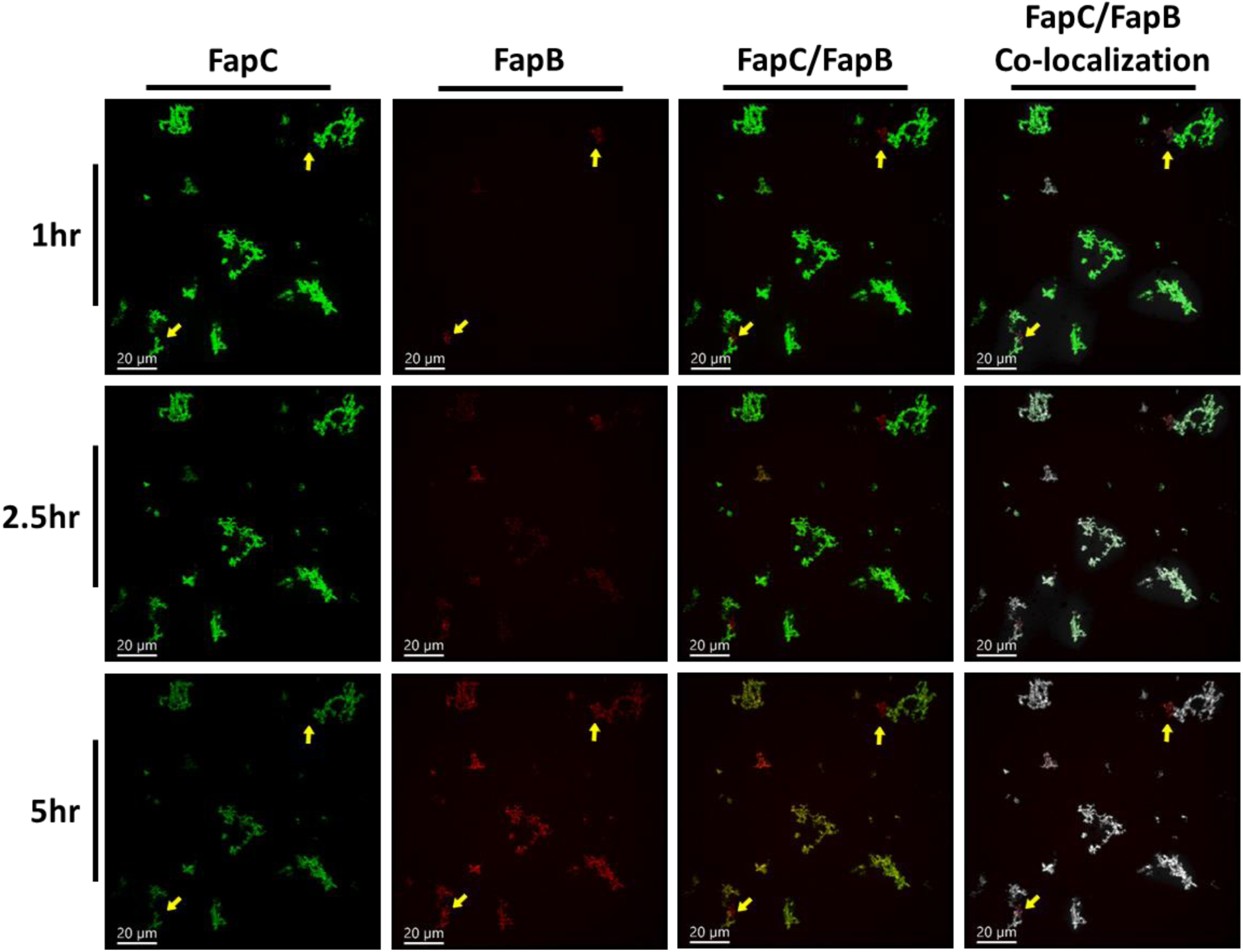
A dynamic visualization of FapB and FapC pre-incubated samples co-localizing over time. Fluorescence microscopy images, captured from [Video 2](https://youtu.be/PPE1Kngsw3U), with FapB and FapC distinctly labeled with Cy3 (red) and Cy5 (green) fluorophores, respectively. The temporal progression of their co-localization is captured at 1, 2.5, and 5 hours, displayed in a sequential order from top to bottom. The merging of Cy3 and Cy5 channels manifests as a yellow/orange hue, marking the regions where FapB and FapC aggregates intersect. To enhance clarity, the right-hand side of each panel row accentuates these co-localization zones in white, as delineated by computational analysis. Yellow arrows are used to pinpoint aggregates of preformed FapB, highlighting their minimal colocalization with FapC over the observed time frame. Incubation during the time-lapse imaging was conducted at pH 7. Scale bars, consistently included in all images, represent a length of 20 μm.

Over a 20-hour time-lapse imaging period, we monitored the formation and localization of aggregates from a 50 µM mixed solution of FapB and FapC at pH 7. Within the first hour, significant FapC aggregates formed, some reaching up to 15µm, while FapB aggregates were smaller. This observation is consistent with kinetic and TEM assays, which indicated a slower fibrillation rate for FapB compared to FapC, especially at pH 7. After 10 hours, there was a notable increase in FapB aggregate formation, primarily co-localized with FapC, intensifying by the 15th hour (Figure 5 and [Video 1](https://youtu.be/U-8s0u023T4)). The quantification of co-localization is detailed in Table S2. The current resolution limits our ability to differentiate between the formation of mixed hetero-fibrils and entangled individual protofilaments.

In our second experimental setup, we explored the dynamics of preformed Fap fibrils, which had been incubated individually for 24 hours, by mixing them together for a 6-hour time-lapse imaging period (Figure 6, [Video 2](https://youtu.be/PPE1Kngsw3U), and Table S2). Initially, our observations revealed a predominance of FapC aggregates and a lesser quantity of FapB aggregates (indicated by yellow arrows in Figure 6), with minimal co-localization between them. As the experiment progressed, we noted a notable increase in the co-localization of FapB aggregates with the pre-existing FapC aggregates. Interestingly, the FapB aggregates that had formed prior to the mixing did not show co-localization with the newly formed FapC aggregates. This observation underscores an asymmetrical interaction pattern between FapB and FapC, where FapC appears to have a more pronounced influence on the behavior of FapB, rather than the other way around. This asymmetry in Fap interaction suggests a directional influence in their aggregation behavior, with FapC playing a more dominant role.

To mitigate the potential impact of photobleaching of the Cy5 dye on FapC amyloids, control samples were prepared under identical conditions but only visualized at the endpoint. These controls exhibited a similar co-localization pattern in both monomer and pre-formed fibril experiments (Figure S6).

### FapB and FapC fibrils exhibit thermostability under extreme heat conditions

To delve deeper into the characteristics of the Fap proteins, we conducted an experiment to evaluate their thermal stability. FapB and FapC were incubated at 37°C for 4 days at pH 5 and 7, respectively, to form fibrils. These fibrils were then exposed to a heat shock at 85°C for 6 and 24 hours. The residual monomers were quantified using SDS-PAGE, taking advantage of the fibril resistance to both SDS buffer and boiling^8,11,42^ (Figure S7). As expected, the 4-day incubation period resulted in a significant reduction in the quantity of monomers. Post heat shock treatment for both 6 and 24 hours, no monomers were detected in either FapB or FapC samples, indicating further fibrillation.

The experiment was escalated to more extreme conditions by autoclaving the pre-formed fibrils for 20 minutes at 121°C. SDS-PAGE analysis revealed the complete disappearance of the monomeric fraction, suggesting additional fibrillation under these harsh conditions. One exemplary SDS-PAGE is presented in Figure S8, and quantification of band intensity with statistical values based on three different experiments is presented in Figure 7A. Incubation alone showed that FapC have fewer residual monomers after four days compared to FapB, confirming a more robust fibrillation. The reduction in residual monomers of both Faps following incubation and more so following autoclaving was consistent, whether the fibrils were grown separately or as a mixture of FapB and FapC. TEM micrographs (Figure 7B) confirmed the thermostability of the fibrils, showing large fibrils intact even after autoclave treatment. These results highlight the extraordinary thermostability of Fap fibrils, suggesting that autoclaving, while likely effective in killing live bacterial cells, may not be sufficient to eliminate the Fap fibrils.

**Figure 7.**
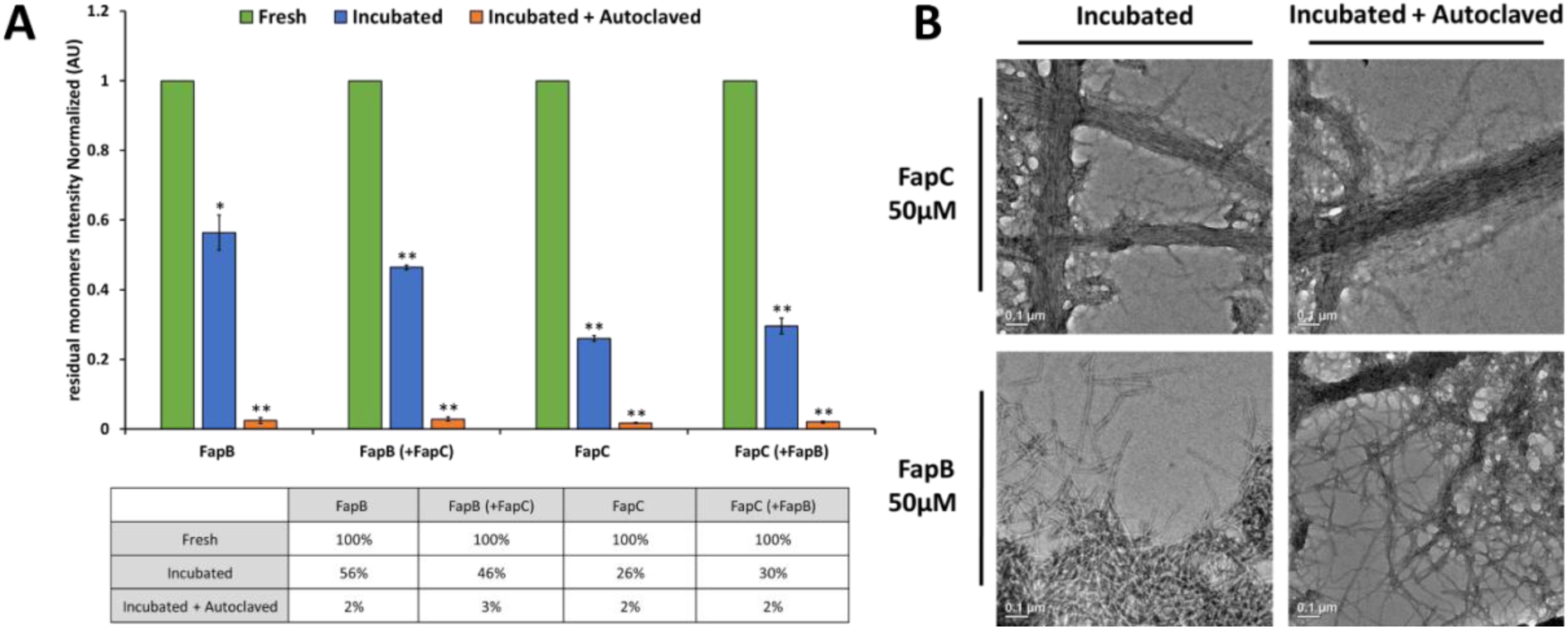
Resilience of FapB and FapC fibril formation under extreme high-temperature autoclaving conditions. (A) The graph presents the quantified band intensities of FapB (incubated at pH 5) and FapC (at pH 7) monomers from SDS-PAGE, averaged over three experimental repetitions (one of which is shown in Figure S8). It compares the residual monomers in fresh samples (green columns), after 4 days of incubation (blue columns), and following both incubation and autoclave treatment (orange columns). The numerical values from the graph are detailed in an accompanying table. Statistical significance was assessed using a T-test (refer to the methods section for details), comparing the incubated and autoclaved samples against the fresh samples. Asterisks indicate levels of statistical significance, with * denoting p<0.05 and ** indicating p<0.001. Error bars represent the standard deviation from the average of the three experimental repetitions. (B) The TEM micrographs display FapB and FapC after 4 days of incubation, both before and after autoclave treatment. These images reveal dense, large fibrils, underscoring the robustness of these structures. The scale bars in all micrographs are set at 0.1 μm.

### Chemical stability of FapB and FapC fibrils

The chemical stability of FapB and FapC fibrils was investigated, given the known resilience of amyloid fibrils to various chemicals like SDS, urea, and guanidinium chloride^42^. Notably, both CsgA and FapC fibrils can be dissolved into monomers by high concentrations of formic acid (FA)^4,42–44^. We used increasing concentrations of FA to assess the stability of FapB and FapC fibrils, incubated individually and together for nine days. The analysis involved SDS-PAGE to detect any residual monomers (Figure 8A), with quantification based on band intensity normalized against fresh, pre-incubated control samples (Figure 8B).

**Figure 8.**
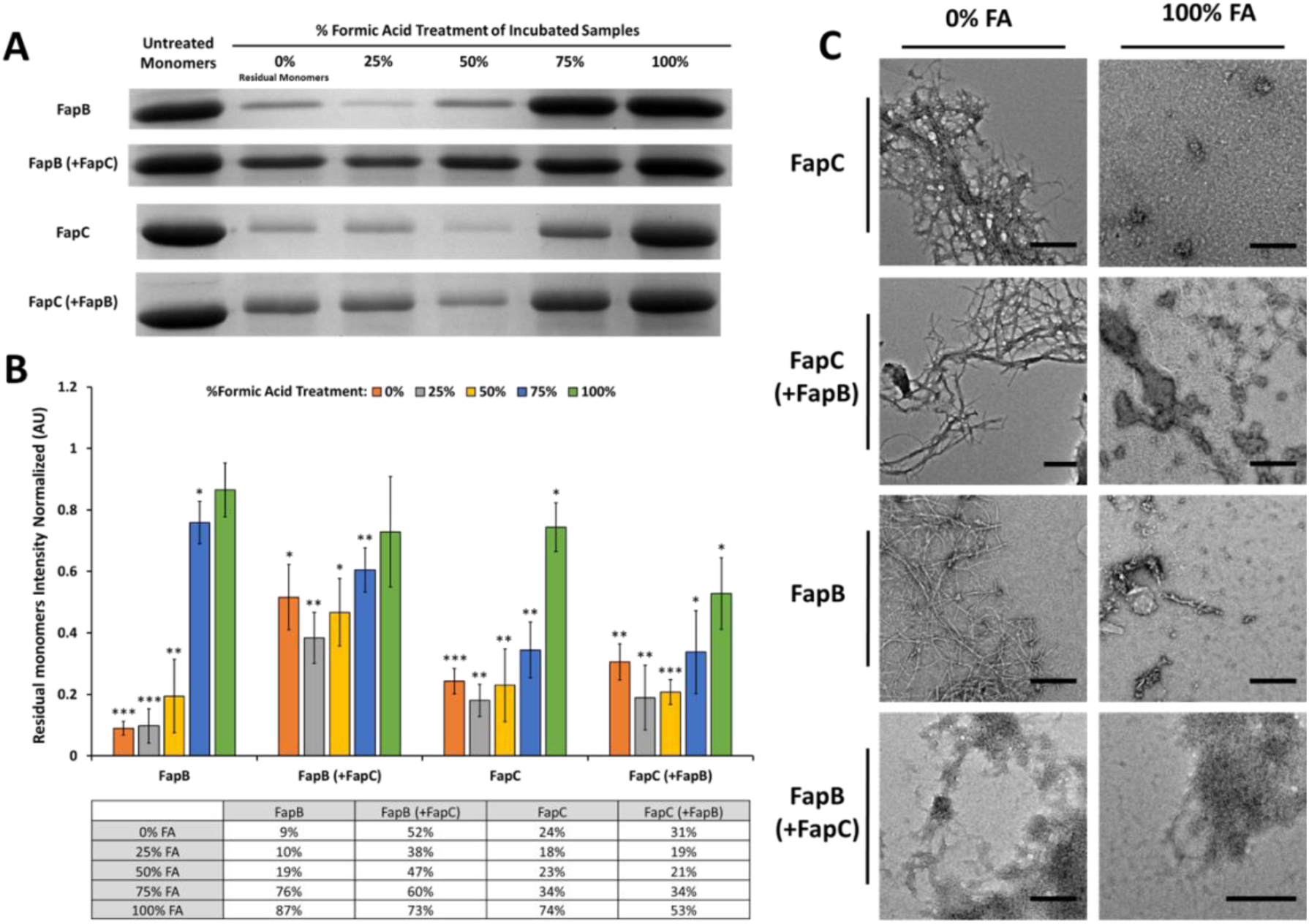
The stability of FapB and FapC fibrils in the presence of formic acid. (A) The figure includes representative SDS-PAGE results of FapB at pH 5 and FapC at pH 7 after a 9-day incubation period, either alone or in combination. This is followed by exposure to escalating concentrations of FA. (B) The quantification of band intensity, derived from three distinct experiments including the one shown in panel A, displays the normalized fraction of soluble monomers in each sample relative to their respective pre-incubated controls. The numerical values from the graph are detailed in an accompanying table. Error bars indicate the standard deviation from the average of the three separate experiments, each conducted on different days. The significance of the differences in band intensity between each sample and the control pre-incubated monomers was assessed using a T-test. Asterisks denote levels of statistical significance: * for p<0.05, ** for p<0.01, and *** for p<0.001. (C) TEM micrographs showcase FapB and FapC after a 9-day incubation, either alone or in combination, followed by treatment with either 0% or 100% FA. The scale bar, applicable to all micrographs, is set at 0.4 μm.

FapB fibrils demonstrated a notable resistance to FA up to a concentration of 50%, with no significant change in the percentage of monomers. However, at 75% FA, a marked disaggregation was observed, resulting in 76% of residual monomers compared to the control pre-incubated samples, a significant increase from the 9% observed without FA. This disaggregation further escalated at undiluted FA, with 87% of the control monomers present. TEM images of FapB confirmed abundant fibril formation after incubation and then significant disaggregation at undiluted FA (Figure 8C). When FapB was co-incubated with FapC, they predominantly formed amorphous aggregates, even in the absence of FA (Figure 8C), and exhibited greater solubility compared to FapB incubated alone, with 52% residual monomers. The addition of FA did not significantly affect the percentage of remaining monomers, with residual monomers ranging up to 73% (Figure 8B). These findings corroborate the inhibitory effect of FapC on the fibrillation of FapB, as observed in the kinetic assays (Figure 3).

FapC displayed remarkable chemical stability, maintaining 34% residual monomers even after the addition of 75% FA, compared to 24% without FA (Figure 8B). With undiluted FA, the soluble fraction was only 74%, indicating the presence of some still resistant fibrils. Interestingly, the presence of FapB did not significantly affect FapC’s stability, with similar percentages of monomers with and without FapB up to 75% FA. Yet, at undiluted FA, the soluble fraction of FapC with FapB was only 53%. TEM micrographs showed thin fibrils of FapC with FapB at pH 7, even with undiluted FA (Figure 8C), supporting a stabilizing effect of FapB on FapC fibrils at this extreme condition.

Comparatively, the mutual effects of FapB and FapC were contrasted with those of curli amyloids CsgA and CsgB (Figure S9). Co-incubation of CsgA and CsgB did not inhibit fibrillation, aligning with CsgB’s role as a nucleator for CsgA. However, their mixed fibrils were less chemically stable than those formed separately, as indicated by higher monomer counts at lower FA percentages. These observations highlight fundamental differences between the curli and Fap systems in terms of the interactions and mutual effects between their amyloid components.

## Discussion

### Differential charge and pH-dependence of Fap amyloids: Implications for environmental adaptation

The distinct charge characteristics and pH sensitivities of FapB and FapC fibrillation highlight their potential roles in environmental adaptability. Drawing parallels with amyloid-β in Alzheimer’s disease, previous studies have shown how environmental factors like pH, ionic strength, temperature, and polypeptide concentration, along with inter-amyloid interactions, critically influence amyloid functions, pathologies, and structures^13,45–48^. The unique pH sensitivities of FapB and FapC underscore their varied interactions with environmental conditions, marked by differences in net charges, isoelectric points, and conservation across orthologs (Table S1 and Figure 2). Their fibrillation responses to pH shifts in PAO1 tested here further emphasize these distinctions (Figures 1&S3).

This diversity in physical and chemical properties among the homologs of FapB and FapC across different *Pseudomonas* strains highlights the complexity and variability of these biofilm-forming proteins. The fibrillation of FapB, which is sensitive to pH and driven by aggregation around its isoelectric point, may even serve as an indicator of environmental changes, and perhaps a preference for different living conditions among various strains. For instance, FapB from *P. aeruginosa* PAO1 and *P. putida* F1 shows a transition from positive to negative net charge around pH 5, whereas FapB from *Pseudomonas* sp. UK4 and *P. fluorescens* Pf-5 only undergoes this switch at pH levels above 9 (Figure 2). This characteristic could be key in regulating fibrillation in vastly different environmental conditions. On the other hand, FapC appears to function as a more robust amyloid, capable of fibrillating across a wide pH range. Its fibrillation process is driven by specific electrostatic interactions that are dependent on its net negative charge. This suggests that the ability of FapC to form fibrils is less influenced by pH changes compared to FapB. Such charge and pH sensitivities, along with higher stability, might represent adaptive strategies, enabling these proteins to function effectively in varied conditions within bacterial biofilms.

*Pseudomonas* species, especially *aeruginosa*, thrive in diverse environments like soil, water, plants, animals, and sewage, which vary significantly in pH, temperature, and salinity^4^. For instance, environmental pH can range from 4.5 to 9.5, demanding substantial bacterial adaptability^4,49^. The optimal conditions for FapC PAO1 fibrillation (pH 6-7) align with the environments in lungs and airways, particularly in cystic fibrosis patients^4,50,51^. Moreover, the more alkaline environment in chronic wounds may favor FapC fibrillation^4,50,51^. Conversely, the preferred conditions for FapB PAO1 fibrillation (pH 5) correlate with the acidity in severely inflamed tissues^52^, suggesting a distinct role for FapB in acute infections and immune responses. Furthermore, the charge differences between FapB and FapC under various conditions might facilitate their interaction. For instance, between pH 5 and 6 in PAO1, FapB could be positively charged while FapC is negatively charged, suggesting potential charge-based interactions during oligomerization and fibrillation.

### Exceptional thermal and chemical resilience of FapB and FapC fibrils

The FapB and FapC fibrils demonstrate remarkable thermal stability, maintaining their fibrous structure even under severe conditions such as autoclaving. Moreover, autoclaving appears to enhance the incorporation of residual monomers into the fibril structure (Figures 7 and S8). This resilience of amyloid fibrils to autoclaving, a process typically lethal to bacteria, has significant implications in both medical and industrial settings. These enduring proteins can adhere to medical instruments and various surfaces, potentially fostering environments favorable for the growth of new bacterial colonies. The presence of these residual amyloids from previous biofilms could facilitate the rapid colonization of surfaces that were initially bacteria-free. This phenomenon raises concerns about the effectiveness of autoclaving in completely eradicating biofilms, highlighting the necessity for a deeper understanding of how these remaining fibrils might promote subsequent bacterial colonization.

Chemically, Fap fibrils demonstrate remarkable stability, resisting disaggregation even under exposure to formic acid. Interestingly, when FapB is present, FapC fibrils exhibit enhanced resilience, with nearly half of the sample remaining in insoluble forms, even in the presence of concentrated, undiluted formic acid (Figure 8). This is in stark contrast to the curli amyloids CsgA and CsgB, which show varying interaction dynamics; mixed fibrils of these proteins are less chemically stable than their individual counterparts (Figure S9).

### Reevaluating the Roles of FapB and FapC in *P. aeruginosa PAO1*

Bacterial functional amyloids, such as Fap proteins, undergo a highly regulated polymerization process, providing physiological benefits to bacteria^18^. Nucleation, a pivotal aspect of this process, influences the formation, growth, kinetics, stability, and morphology of amyloid fibrils, along with their physiological and pathological role ^13,18,53^. In *P. aeruginosa*, FapB has been hypothesized to initiate FapC fibrillation, similar to CsgB’s role, considering the lower abundance of FapB in biofilms ^4,8,11,25,54^. However, our recent findings challenge this perspective.

When FapB and FapC from PAO1 are combined, we observed a prolonged lag phase in fibrillation (Figure 3), indicating a delay in β-rich fibril formation. Additionally, SDS-PAGE analysis of remaining monomers revealed that in each other’s presence, Fap proteins maintain higher solubility over nine days compared to when isolated (Figures 3&8). FapB’s aggregation rate is more influenced by FapC than vice versa. Moreover, FapB fibril seeds did not significantly alter the fibrillation rate of FapC, whereas FapC seeds modestly enhanced FapB fibrillation (Figure 4). Confocal microscopy supports this, showing FapB aggregation predominantly colocalize with pre-existing FapC aggregates, while the reverse is less common (Figures 5&6 and Videos S1&S2). Both Faps showed slightly increased fibrillation with self-seeding (Figure 4), aligning with patterns observed in other amyloids^55–58^.

These findings suggest that FapB and FapC employ distinct nucleation and growth strategies, with their interaction in combined amyloid formations being complex and condition-dependent. This raises questions about the role of FapB as a primary nucleator for FapC, at least in the PAO1 orthologs, hinting at a more intricate relationship. The co-localization of Faps, facilitated by interactions between their monomers or fibrils, might lead to mixed fibrils or intertwined protofilaments, potentially enhancing the structural integrity of *P. aeruginosa* biofilms in diverse environments. Given FapC faster fibrillation rate and reduced pH-dependency, it appears to be the primary amyloid for biofilm scaffolding, similar to CsgA. In contrast, FapB might serve as an auxiliary amyloid, fibrillating under specific conditions and integrating into the biofilm through self-nucleation or existing FapC fibrils. Moreover, the complex dynamics within the Fap system could be influenced by other proteins like FapA and FapE. FapA is known to modulate FapC fibrillation and alter FapB’s distribution in mature fibrils^8,9,20,54^, while FapE acts as an extracellular chaperone during fibrillation and is present within mature amyloid fibrils^8,24,54^. This intricate interplay of proteins within the Fap system exemplifies the sophisticated regulation of amyloid interactions in bacterial biofilms.

### Conclusions

Given the significant health and environmental impacts of *Pseudomonas* infections, particularly the resistance of *P. aeruginosa* biofilms to antimicrobial treatments and their ability to evade host defenses^59–64^, a deeper understanding of their virulent amyloids is essential. Our research uncovers the remarkable thermal and chemical resilience of FapB and FapC fibrils. This resilience indicates that biofilm structures may withstand standard sterilization processes, potentially contributing to the persistence of *P. aeruginosa* infections. The durability of these fibrils also presents exciting opportunities for developing robust bio-nanomaterials or protective coatings, capitalizing on their unique properties.

Our studies also illuminate the significant influence of environmental factors on Fap proteins, suggesting that homologs in different *Pseudomonas* strains may have adapted to specific environments and roles. Challenging the view of FapB as the primary nucleator, our findings indicate a more pronounced role of FapC in influencing FapB aggregation. However, the interplay between FapB and FapC is intricate: FapB aids in stabilizing FapC fibrils, while FapC can slow down the fibrillation rate of FapB, yet still serve as a cross-seeding template. This complex dynamic is key to understanding their roles within bacterial biofilms. Furthermore, the observed distinctions between Fap and curli amyloids underscore the importance of comprehensively understanding different amyloid systems for the development of effective intervention strategies.

## Methods

### Materials and Reagents

Competent E. coli BL-21 cells (New England Biolabs) were used for the transformation of pET-28b(+) plasmids (Novagen). Phenylmethylsulfonyl fluoride protease inhibitor (Sigma-Aldrich); guanidinium HCl (SPECTRUM) were employed for the lysis buffer during protein purification. Isopropyl β-D-1-thiogalactopyranoside (IPTG) purchased from Ornat was used to induce protein expression. Cyanine3 NHS ester and Cyanine5 NHS ester fluorophore dyes were purchased from Lumiprobe and dissolved in DMSO (Merck). 15µ-Slide 8 Well Glass Bottom/15µ-Slide VI 0.4 were purchased from ibidi GmbH. HisPur cobalt resin beads (Thermo Scientific via Danyel Biotech) were used for protein purification. Luria-Bertani (LB) agar, Bacto Yeast, Bacto Tryptone (BD Biosciences), and Kanamycin antibiotic (MERCK group) were used for bacteria growing. GelRed Nucleic Acid Stain (Gold Biotechnology) was used for DNA gel staining. Formic acid was purchased from Gadot Group. Ultra-pure water (Biological Industries) were used throughout the study. Thioflavin T (ThT) dye was purchased from Sigma-Aldrich. Acrylamide/bis-acrylamide 29:1 40% (BioLab), Ammonium Peroxydisulfate (Alfa Aesar) and N, N, N’, N’-Tetramethylethylenediamine (Sigma Aldrich) were used for gel preparation. Sodium Dodecyl Sulfate (Holland Moran group), DL Dithiothreitol (Sigma Aldrich) and ExcelBand™ 3-color Regular Range Protein Marker (SMOBIO) were used in SDS-PAGE analysis. InstantBlue® Coomassie Protein Stain from Expedeon was used for protein gel staining. 400 mesh copper grids with support films Formvar/Carbon (Ted Pella Inc) and 1% uranyl acetate solution (Electron Microscopy Sciences Ltd) were used for negative staining in transmission electron microscopy (TEM). Thermal seal film (EXCEL scientific), and black 96-well flat-bottom plates (Greiner bio-one) were used for kinetic assay measurements. Zeba spin desalting columns (Rhenium) and disposable polypropylene columns (Bio-rad) were used for buffer exchange and purification steps. Potassium dihydrogen phosphate (Merck Group), Potassium phosphate dibasic (SPECTRUM), sodium acetate (Merck Group), Tris Base (Fisher Bioreagents), Bis-Tris Hydrochloride (Sigma Aldrich), and sodium chloride (BioLab) were used for buffer preparation.

### Computational analyses and calculations of biophysical properties

To explore the biophysical attributes of FapB and FapC amyloids, we conducted a detailed computational analysis, focusing on their length, pH sensitivity, and charge characteristics. This investigation encompassed amyloid homologs from various *Pseudomonas* strains: *P. aeruginosa* PAO1 (UniProt# Q9I2E9, Q9I2F0), *P. fluorescens* Pf-5 (UniProt# Q4KC08, Q4KC07), *P. putida* F1 (UniProt# A5W4A4, A5W4A5), and *Pseudomonas* sp. UK4 (UniProt# C4IN69, C4IN70). We utilized the UniProt database and the Prot-pi web calculator (https://www.protpi.ch/Calculator/ProteinTool) for precise sequence analysis of each homolog, including length, charge, and isoelectric point. Prot-pi was also used to calculate the overall charge of each amyloid at different pH levels (4 to 9). For FapB and FapC from the *P. aeruginosa* PAO1 strain, used in our experiments, we calculated the overall charge at various pH levels also considering the impact of a Histidine tag at the N- or C-terminal site.

Molecular insights into the protonation state of FapB under various pH conditions were obtained using the AlphaFold2 artificial intelligence-based structure modeling tool^36,37^. Structure predictions for *P. aeruginosa* PAO1 FapB and its Histidine-tagged variant were carried out on the online Google Colab service using AlphaFold v2.1.0 advanced. To assess the protonation effects on these structures, they were converted to PDB files and analyzed with the H++ automated system^65,66^. This system calculates pK values of ionizable groups in macromolecules and adds hydrogen atoms as needed for the specified environmental pH. The resulting structures, showing electrostatic potential, were visualized, colored, and analyzed using UCSF ChimeraX software^67^, allowing us to compare variations of charge distribution over the protein surface at different pH levels.

The [Clustal Omega Sequence Alignment tool](https://www.ebi.ac.uk/jdispatcher/msa/clustalo)^68,69^ was used to determine the sequence identity between FapB and FapC from *P. aeruginosa* PAO1, focusing on their full sequences, excluding signal peptides. Similarly, we analyzed the sequence identity between E. coli K12’s CsgA and CsgB (UniProt# P28307, P0ABK7).

### Protein production of FapB and FapC

#### Construction of FapB and FapC plasmids

The genes of the PAO1 *Pseudomonas aeruginosa* strain FapB and FapC (UniProt #: Q9I2E9 and Q9I2F0, respectively) were successfully cloned into the pET28b expression vector plasmid. We designed specific primers using the NEB site, ensuring they were complementary to targeted regions of the genes and included necessary restriction sites. These sites were strategically selected to allow simultaneous cutting of both the vector and the gene with the same restriction enzymes, thereby streamlining the gene insertion process.

The NEB Q5® High-Fidelity PCR Kit was utilized to amplify the genes from *P. aeruginosa* genomic DNA. Subsequently, the amplified genes were cloned into the pET28b(+) plasmid (Novagen). Notably, the pET28b vector incorporates a Histidine tag at either the N- or C-terminal site, and the primers were tailored to align with specific segments of the gene sequence. This approach led to the creation of four distinct plasmids: two containing the FapB gene and two containing the FapC gene, each with a Histidine tag positioned at one of the terminal sites.

To confirm the successful integration of the genes within the pET28b vector, several steps were undertaken. Initially, the plasmids were transformed into *E. coli* XL-1 competent cells and cultured on LB agar plates supplemented with 50 μg/ml kanamycin antibiotic. Following overnight incubation at 37°C, selected colonies were propagated in LB medium. The plasmids were then extracted using the Presto Mini Plasmid Kit, and their concentrations were quantified with a NanoDrop spectrophotometer.

To ascertain the presence of the correct genes in the plasmids, we employed specific restriction enzymes (XhoI, NcoI, and NdeI from NEB) to cleave the plasmids at predetermined sites corresponding to the gene insertion. The resultant plasmid fragments were separated via electrophoresis on a 1% agarose DNA gel, which was stained with GelRed Nucleic Acid Stain. The gel documentation was performed using the Bio-Rad Gel imaging system to verify the precise insertion of the genes into each plasmid. Finally, to authenticate the gene sequences, DNA sequencing was conducted by Macrogen - Syntezza Bioscience Ltd, Jerusalem, ensuring the accuracy and integrity of our plasmid constructs.

#### Expression and purification of FapB and FapC

The production of FapB and FapC proteins was carried out separately using the following protocol. All buffers were filtered through a 0.22 μm Millex filter and maintained at 4°C. The purified pET28b plasmids, containing either the FapB or FapC genes with a Histidine tag at the N- or C-terminal site, were transformed into competent *E. coli* BL-21 cells. These were then plated on LB agar plates supplemented with 50 μg/ml kanamycin antibiotic and incubated overnight at 37°C. Colonies from these plates were transferred to 100 ml LB medium containing the antibiotic and grown overnight at 37°C.

The overnight cultures were diluted into 700 ml of the same medium and incubated at 37°C with shaking at 220 rpm until an optical density (OD) of 0.8-0.9 at 600 nm was reached. Protein expression was induced by adding 1 mM isopropyl β-D-1-thiogalactopyranoside (IPTG), followed by a 3-hour incubation under the same conditions. The bacterial cells were harvested by centrifugation at 4,500 rpm for 30 minutes and stored at -80°C.

*Protein Purification*: For purification, HisPur cobalt resin beads (Thermo Scientific) were used. The cell pellets were thawed, resuspended in 30 ml of lysis buffer (6 M guanidinium HCl, 50 mM potassium phosphate, 0.1 mM PMSF, pH 7.4), and incubated overnight at 4°C with agitation. The lysate was centrifuged at 17,000g for 30 minutes at 4°C to separate into supernatant and pellet fractions. The supernatant was incubated with 2 ml of equilibrated HisPur cobalt resin beads in lysis buffer for 2-3 hours at 4°C with agitation. The mixture was then loaded onto a disposable polypropylene column at 4°C and washed with 20 ml of 50 mM potassium phosphate buffer (pH 7.4), followed by a wash with the same buffer containing 10 mM imidazole. Proteins were eluted using 350 mM imidazole in 50 mM potassium phosphate buffer (pH 7.4). Imidazole was removed, and the buffer was exchanged using Zeba spin desalting columns (7k mwco, ThermoFisher Scientific) at 4°C into different buffers as required for specific experiments.

*Protein concentration and verification*: Protein concentrations were determined by measuring absorbance at 280 nm using extinction coefficients of 24,535 M^-1^ cm^-1^ for FapC and 1,490 M^-1^ cm^-1^ for FapB, calculated using the Expasy server. The identity of FapB and FapC proteins was confirmed through Western blot analysis using an anti-Histidine tag antibody.

### Purification and labeling of FapB and FapC with fluorophores

To observe the formation of FapB and FapC fibrils, we utilized two fluorophore dyes: Cyanine3 NHS ester (Excitation: 555 nm, Emission: 570 nm) and Cyanine5 NHS ester (Excitation: 646 nm, Emission: 662 nm) from Lumiprobe. These NHS ester dyes covalently bond with primary amines, predominantly lysine, in proteins, ensuring stable labeling. Each fluorophore was dissolved in DMSO to achieve a concentration of 4 mg/ml. Protein lysate samples were split into two portions. One served as an unlabeled control, while the other was designated for labeling.

*Labeling Process*: The supernatant fraction was mixed with 1 ml of HisPur cobalt resin beads pre-equilibrated with lysis buffer and agitated for 1-2 hours at 4°C. After loading the mixture onto a disposable polypropylene column, it was washed with 10 ml of 50 mM potassium phosphate buffer, followed by a wash with the same buffer containing 10 mM imidazole. A 2:1 molar ratio of dye to protein was prepared for either Cy3 or Cy5 NHS ester, using the concentration of the unlabeled fraction as a reference. The dye solution was then mixed with the HisPur-bound amyloid samples (FapB with Cy3 and FapC with Cy5) and incubated on the column for 2 hours at 4°C in the dark. Additional washes were performed, including a quenching buffer (50 mM Tris-HCl, pH 7.4), followed by two washes with 50 mM potassium phosphate buffer. The labeled amyloids were eluted, and imidazole was removed as previously described. The protein aliquots were stored at -80°C.

*Concentration determination and verification*: To quantify the fluorophore-tagged FapB and FapC, SDS-PAGE analysis was conducted. Tagged and untagged protein samples were heated at 95°C for 10 minutes with a sample buffer containing SDS and DTT, then loaded onto 15% polyacrylamide gels and run at 120V for 60 minutes. A Typhoon™ FLA 70000 Phosphorimager was used to confirm correct labeling, followed by staining with InstantBlue® Coomassie Protein Stain for 30 minutes. Gel imaging was performed using a BioRad gel doc system, and the concentrations of the fluorophore-dyed proteins were determined by comparing band intensities to those of the known concentrations of the untagged proteins.

### Fluorescence microscopy analysis of FapB and FapC

For the fluorescence microscopy studies, the labeled FapB and FapC protein samples were first thawed. They were either mixed in a 1:1 ratio to achieve a concentration of 50 µM in 50 mM potassium phosphate buffer or incubated separately for 24 hours at 37 °C in the dark before mixing. Both the mixed samples and the unmixed controls were then placed onto 8-well chamber slides (15µ-slides 8-well glass bottom - Ibidi) for observation under a confocal microscope. The imaging was performed using a Nikon Ti2-Eclipse microscope, equipped with a 100x/1.35 silicone immersion objective (Nikon CFI SR HP Plan Apochromat Lambda S) and 561nm and 640nm laser diodes. Photometrics BSI sCMOS cameras were used for capturing the images. The imaging was focused on the bottom of the well showing residing aggregates.

*Co-localization analysis*: To analyze the co-localization of FapB (labeled with Cy3) and FapC (labeled with Cy5), we utilized 6-μm-thick z-stacks imaged at 100x magnification. Time-lapse images were recorded over approximately 20 hours and 6 hours in two separate experiments (one with monomeric proteins and the other with preincubated fibrils). Photobleaching correction was applied to these images using the Napari plugin with a bi-exponential method. The data analysis was conducted using Imaris 10.1 software (Oxford Instruments). The Coloc tool in Imaris was employed to identify co-localization areas, creating a colocalization channel based on a fixed minimal intensity threshold. For quantitative analysis, images were processed in Imaris to generate 3D surfaces for each channel, again based on a fixed intensity threshold. We measured the average total volume, the percentage of average overlapped volume relative to the total surface, and the total number of voxels for each channel using Imaris’ inbuilt functions. The detailed data can be found in Table S2.

### Thioflavin T fluorescence kinetic assays for monitoring fibrillation of FapB and FapC

Thioflavin T (ThT) assays are a standard method for studying amyloid fibril formation. We used this approach to investigate the fibrillation kinetics of FapB and FapC under various conditions, including different pH levels, ionic strengths, co-incubation ratios, and the presence of preformed fibril seeds.

#### ThT assay of Faps at varied pH and ionic strength conditions

Freshly purified monomers of FapB and FapC were transitioned from the elution buffer to the desired condition buffer using Zeba spin desalting columns. Based on preliminary assessments, we selected N-terminal tagged FapB for its higher fibrillation propensity (Figure S2) and C-terminal tagged FapC, as the tag’s position showed no significant impact on its fibrillation. Initial fibrillation tests were conducted in a 50 mM potassium phosphate buffer at pH 7.4. To explore the effect of pH, we employed a “Universal Buffer” covering a pH range of 2-10^70^, consisting of 20 μM Tris, 20 μM Bis-Tris, and 20 μM Sodium acetate. ThT assays were performed by combining FapB or FapC monomers at concentrations of 25, 50, and 100 μM with 20 μM ThT prepared as a stock solution in ultra-pure water and diluted in the respective pH buffer, in a total volume of 100 μl. As controls, wells containing only 20 μM ThT were included for each pH value examined. Additionally, to assess the influence of ionic strength on fibrillation, we used a pH 7 buffer supplemented with NaCl to achieve final concentrations of 0, 0.15, and 0.3 M NaCl. The reaction mixtures were placed in black 96-well flat-bottom plates (Greiner bio-one) covered with thermal seal film (EXCEL scientific), and incubated in a plate reader (FLUOstar Omega, BMG Labtech) at 37°C, with orbital shaking at 300 rpm for 30 seconds prior to each measurement. ThT fluorescence was recorded every 3 minutes, utilizing an excitation wavelength of 438 ± 20 nm and an emission wavelength of 490 ± 20 nm. All measurements were performed in triplicate, and the entire experiment was repeated at least three times.

#### ThT assays of Faps at different co-incubation ratios

To investigate the impact of interaction/nucleation between FapB and FapC monomers on their fibrillation kinetics, we mixed freshly purified FapB monomers with increasing concentrations of FapC monomers in the “universal buffer” at pH 5, along with filtered ThT prepared in ultra-pure water. The final concentrations in the reaction wells were 50 μM FapB mixed with 0, 2, 5, 10, 25, 50, 75, or 100 μM FapC, along with 20 μM ThT, in a final volume of 100 μl. Control wells contained only FapC at each concentration and ThT, replacing FapB with an equivalent volume of the “universal buffer” at pH 5. A reciprocal experiment was conducted by mixing FapC monomers with increasing concentrations of FapB monomers at pH 7, following the same protocol with the same concentrations. In this case, the control wells exclusively contained FapB and ThT at pH 7. The measurement settings were consistent with those described in the previous paragraph. The measurements themselves were performed in triplicate, and the entire experiment was repeated at least three times.

#### ThT assays of self- and cross-seeding of the Faps

The interaction between FapB and FapC was further investigated by assessing the influence of pre-formed sonicated fibrils, referred to as “seeds,” on the fibrillation rate of their respective monomers using a ThT kinetic assay. For this purpose, we incubated 50 μM of FapB or FapC with varying volumes of pre-formed sonicated fibrils of FapB and FapC. To produce FapB and FapC fibrils, freshly purified monomers were incubated in a final volume of 350 μl and a concentration of 100 μM in a “universal buffer” at pH 5 or 7 (respectively) for one week at 37°C with shaking at 300 rpm. The fibrils were then dried under a vacuum for 24 hours, followed by re-suspension in their original buffer and vigorous shaking. Subsequently, the fibrils were bath sonicated at room temperature for 5 minutes. For further sonication, the fibrils were placed on ice and subjected to VC750 VibraCell (Sonics, CT, USA) tip sonication at 30% amplitude, with three 20-second bursts at 50-second intervals.

To perform the measurements, freshly purified FapB monomers were mixed with increasing volumes of pre-formed sonicated fibrils of FapB and FapC (self-seeding and cross-seeding) in the same buffer at pH 5. Similarly, FapC monomers were mixed with the corresponding pre-formed sonicated fibrils of FapB and FapC, with the only difference being the pH set to 7. The final concentrations in the reaction wells were 50 μM FapB or FapC monomers mixed with 0, 2, 5, and 10% (v/v) of seeds, along with 20 μM ThT, in a final volume of 100 μl. Controls included wells containing only the seeds at each concentration, along with ThT and the respective pH. The measurement settings remained consistent with those described in the previous paragraph. All measurements were performed in triplicate, and the entire experiment was repeated at least three times.

### Transmission Electron Microscopy

Transmission Electron Microscopy (TEM) was utilized to visualize the fibrils of FapB and FapC, providing insights into the effects of various fibrillation conditions and treatments on their structural formation. Samples for TEM analysis were collected either at the conclusion of the ThT kinetic assays or at predetermined intervals based on the assay’s protocol. In instances where the fibrils underwent specific treatments, such as temperature variations or exposure to different concentrations of formic acid, samples were taken directly from these treated conditions.

For TEM grid preparation, a volume of 5 μl from the sample was applied onto 400 mesh copper grids. These grids were equipped with support films made of Formvar/Carbon, obtained from Ted Pella, and were charged using high-voltage, alternating current glow-discharge immediately prior to sample application. This preparation ensured optimal adherence of the samples to the grids. After allowing the samples to settle on the grids for 2 minutes, they were subjected to negative staining using a 1% uranyl acetate solution for an additional 2 minutes.

The imaging of the samples was conducted using an FEI Tecnai G2 T20 S-Twin transmission electron microscope. This microscope operated at an accelerating voltage of 200 KeV, enabling high-resolution visualization of the fibril structures. The micrographs were captured at the MIKA Electron Microscopy Center, which is part of the Department of Material Science & Engineering at the Technion – Israel Institute of Technology.

### SDS-PAGE analysis of FapB and FapC thermostability

To assess the stability of FapB and FapC fibrils, we conducted SDS-PAGE analysis to quantify residual monomers, considering that insoluble fibrils do not migrate in SDS-PAGE even after boiling. This characteristic is shared with other amyloids involved in biofilm formation, such as CsgA^41^.

We investigated the thermostability of FapB and FapC fibrils, as well as mixed fibrils formed by combining both amyloids. Freshly purified FapB and FapC were incubated separately in a “universal buffer” at pH 5 and 7, respectively. For mixed fibril samples, we prepared two sets, one at pH 5 and the other at pH 7. Each amyloid was at a final concentration of 50 μM, and in the mixed samples, they were combined in a 1:1 ratio. These samples were incubated for 4 days at 37°C with shaking at 300 rpm. Following this initial incubation, the samples underwent thermal treatments. One set was incubated at 85°C for 0, 6, and 24 hours, while another set was exposed to extreme temperatures by autoclaving at 121°C. Post-treatment, each sample was mixed with SDS sample buffer supplemented with DTT and heated at 95°C for 10 minutes. To quantify the amount of monomers obtained post-treatment, we collected a sample of fresh monomers before the initial incubation and fibril formation as a control.

The samples were then loaded onto a 15% acrylamide SDS-PAGE gel and subjected to electrophoresis at a constant voltage of 120V for 60 minutes. After electrophoresis, we used Coomassie Stain (Expedeon - InstantBlue Coomassie Protein Stain) to visualize and stain the soluble monomers of FapB and FapC migrating within the gel. Gel imaging and quantification were performed using a Gel Doc–BioRad system. The quantified values reported were calculated by normalizing the intensity of the post-treatment bands to that of a control band, of the untreated monomer sample of the same quantity. This normalization ensures that all numerical values displayed in the graphs and tables represent the percentage of monomers remaining in the samples relative to the initial monomer quantity. The entire experiment was replicated twice for the heat shock treatment and a minimum of three times for the autoclaving process. The quantification of the results from the autoclaving treatment was derived from data across three separate, independent experiments.

### SDS-PAGE analysis of FapB and FapC stability in formic acid

To assess the chemical stability of FapB and FapC amyloid fibrils, we focused on their interaction with formic acid (FA), a solvent known for its ability to disrupt amyloid structures, including dissolving amyloid beta plaques^43,44^. In this experiment, we prepared samples of FapB and FapC separated or mixed, following the methodology outlined in the previous section. However, in this instance, we allowed the samples to undergo fibrillation for an extended period of 7-10 days.

Post-fibrillation, the samples were dried using a SpeedVac (RVC 2–18 CDplus, CHRIST) to eliminate excess liquid and concentrate the fibrils. Subsequently, each dried amyloid sample was reconstituted in varying concentrations of FA, diluted in ultrapure water. The FA concentrations ranged from 0% to 100%, with increments of 25%. We then adjusted the volume of each resuspended sample back to its original 200 μl volume prior to drying. Following resuspension, the samples were incubated for 10 minutes at 25°C and then rapidly frozen using liquid nitrogen. Overnight lyophilization (ScanVac Cool Safe - Freeze dryers, LABOGENE) was employed to remove any remaining liquid from the samples. For the analysis, the lyophilized amyloids were reconstituted to their initial volume and concentration of 50 μM using the appropriate “universal buffer” at either pH 5 or pH 7. The reconstituted samples were then subjected to SDS-PAGE, normalized, and analyzed as previously described. The entire experiment was replicated three times, and quantification and statistics were derived from data across three separate, independent experiments.

### SDS-PAGE analysis of *E. coli* CsgA and CsgB stability in formic acid

#### *E. coli* CsgA and CsgB protein production

Electro-competent BL21-DE3 slyD knockout cells were electroporated with pET11a plasmids carrying the CsgA or CsgB genes. These transformed cells were cultured on LB-agar plates with ampicillin at 37°C. A single colony was used to initiate a preculture, which then started the main expression culture in 2xYT medium supplemented with ampicillin and glucose. This culture was incubated at 37°C with shaking until reaching a specific bacterial density, followed by IPTG induction and further incubation. Post-incubation, the culture was centrifuged, and the cell pellet was stored or prepared for purification.

*Purification*: The cell pellet was dissolved, centrifuged to remove debris, and then incubated with Ni-NTA beads for binding. After several washes and elution steps, the protein concentration was determined using a DeNovix DS-11 Nanodrop. The purified protein was filtered, snap-frozen, and stored. For assays, the protein was thawed, desalted, and its concentration re-assessed.

#### Fibrillation of CsgA and CsgB

Fibrils were generated from protein samples with concentrations of 30 µM. These samples were incubated at 37°C in Eppendorf tubes, either individually or mixed in a 1:1 molar ratio, for 48 hours. Post-incubation, the fibrils underwent a triple washing process with centrifugation at 13,000g for 10 minutes. The supernatant was removed, and the fibril pellet was resuspended in fresh 50 mM Tris buffer (pH 7.4). The protein concentration in the supernatant was quantified. The washed fibrils were sonicated using a Fisherbrand 505 Sonicator.

#### Formic acid stability assay

The sonicated fibril samples were transferred to new Eppendorf tubes and centrifuged at 13,000g for 10 minutes. The supernatant was discarded, and the fibril pellet was treated with 50 µl of X% formic acid, mixed, and vortexed gently. After a 20-minute room temperature incubation, the samples were centrifuged again. A portion of the supernatant was flash-frozen in liquid nitrogen, lyophilized, and reconstituted in 2x Laemmli SDS-PAGE Loading Buffer. The samples were then loaded onto a Bio-Rad TGX Stain-Free 4-20% SDS-PAGE gel and electrophoresed.

Protein bands were detected using Coomassie staining and Stain-Free imaging. While Coomassie staining could not differentiate between CsgA and CsgB, Stain-Free imaging selectively visualized CsgA based on its tryptophan residues. Band intensities were analyzed using Bio-Rad’s Image Lab Software and normalized against those observed in the 98% formic acid sample for comparative analysis.

## Acknowledgments

M.L. acknowledges research support from the Israel Science Foundation, Grants No. 2111/20 the Cure Alzheimer’s Fund, and the Russell Berrie Nanotechnology Institue (RBNI). D.E.O. acknowledges support from the Sino-Danish Center (SDC) and the Lundbeck Foundation (grant no. R276-2018-671). We thank Amit Efroni for support in Faps initial protein production and Eilon Barnea for critical advice and support. We acknowledge guidance from Yaron Kauffmann from the MIKA Electron Microscopy Center of the Department of Material Science & Engineering at the Technion, and Ellina Kesselman from the Center for Electron Microscopy of Soft Matter of Wolfson Department of Chemical Engineering at the Technion, Israel. We acknowledge guidance and support from Nitsan Dahan and Yael Lupu-Haber from the Microscopy core facility center at the Lorry I. Lokey Interdisciplinary Center for Life Sciences and Engineering. We would like to acknowledge the use of OpenAI’s GPT-4 model to improve the quality of writing in this study.

